# Measuring site-specific glycosylation similarity between influenza A virus variants with statistical certainty

**DOI:** 10.1101/2020.03.13.991380

**Authors:** Deborah Chang, William E. Hackett, Lei Zhong, Xiu-Feng Wan, Joseph Zaia

## Abstract

Influenza A virus (IAV) mutates rapidly, resulting in antigenic drift and poor year-to-year vaccine effectiveness. One challenge in designing effective vaccines is that genetic mutations frequently cause amino acid variations in IAV envelope protein hemagglutinin (HA) that create new *N*-glycosylation sequons; resulting *N*-glycans cause antigenic shielding, allowing viral escape from adaptive immune responses. Vaccine candidate strain selection currently involves correlating antigenicity with HA protein sequence among circulating strains, but quantitative comparison of site-specific glycosylation information may likely improve the ability to design vaccines with broader effectiveness against evolving strains. However, there is poor understanding of the influence of glycosylation on immunodominance, antigenicity, and immunogenicity of HA, and there are no well-tested methods for comparing glycosylation similarity among virus samples. Here, we present a method for statistically rigorous quantification of similarity between two related virus strains that considers the presence and abundance of glycopeptide glycoforms. We demonstrate the strength of our approach by determining that there was a quantifiable difference in glycosylation at the protein level between wild-type IAV HA from A/Switzerland/9715293/2013 (SWZ13) and a mutant strain of SWZ13, even though no *N*-glycosylation sequons were changed. We determined site-specifically that WT and mutant HA have varying similarity at the glycosylation sites of the head domain, reflecting competing pressures to evade host immune response while retaining viral fitness. To our knowledge, our results are the first to quantify changes in glycosylation state that occur in related proteins of considerable glycan heterogeneity. Our results provide a method for understanding how changes in glycosylation state are correlated with variations in protein sequence, which is necessary for improving IAV vaccine strain selection. Understanding glycosylation will be especially important as we find new expression vectors for vaccine production, as glycosylation state depends greatly on the host species.

## Introduction

In the 2017-2018 influenza season, nearly 80,000 influenza-related deaths were reported in the U.S., the highest death toll in 40 years (1). Effective vaccines help to mitigate disease, but influenza A virus (IAV) mutates rapidly, resulting in antigenic drift that causes the need for frequent updates to the influenza seasonal vaccine strains. IAV mutates during viral evolution in two ways. First, antigenic drift causes changes in the protein sequence to accumulate over time, altering the antigenic reactivity of the virus. Such changes may add *N*-glycosylation sequons that cause antigenic shielding, alter receptor binding avidity, and change the interactions with innate immune system lectins (2). Second, antigenic shift occurs when IAV RNA segments recombine with zoonotic genes to create new pandemic strains (3). New pandemic strains start out with very low glycosylation in the head group, reflecting different selective pressures on IAV in animal hosts than in humans (H1N1 re-assorted from swine in 2009).

The antigenic character of the abundant IAV envelope protein hemagglutinin (HA) depends on protein sequence (4, 5) and glycosylation state (6, 7). There are defined antigenic regions of HA protein sequence that evolve rapidly (8). Antigenic regions must balance their ability to evade host adaptive immune response with maintaining receptor-binding function (9). HA antigenicity can be assessed through hemagglutination inhibition (HI) data. Correlating HI with protein sequence information allows prediction of antigenicity based on protein sequence (10–12). HA surface antigenicity is affected strongly by the presence and types of *N*-glycosylation, which cannot be predicted from genetic sequence or 3D protein coordinates (13). *N*-Glycosylation sequons accumulate in the protein sequence, giving rise to increasing glycosylation as the strain evolves, shielding of antigenic sites (8, 14, 15), influencing receptor binding (16, 17) and binding to innate immune system molecules (18, 19).

Previous research demonstrated the power of mass spectrometry to identify HA *N*-glycans site-specifically (20–23). However, most glycosylated HA models in the literature were constructed using unoptimized, generic glycans without defined information about the glycan antennae. Our recent MCP article is an exception (24). *N*-Glycosylation is microheterogeneous—there is a distribution of possible glycans at each site—and certain sites may be inaccessible to processing by ER mannosidases compared to other sites (24). Varying degrees of glycan processing affect the lectin-binding and other host interaction properties at these sites. HA is macroheterogeneous; a population of HA proteoforms exists. The glycosylated proteoforms that are most immunogenic are not known. In order to correlate antigenicity and immunogenicity with glycosylation, rigorous comparison of glycosylation similarities among strains is required (25). Furthermore, the viral expression systems that can produce the most immunodominant glycans are also unknown.

Year-to-year influenza vaccine effectiveness is low, ranging from 10-60% in the years 2004-2018 (26). For effective vaccine design, antigenicity of the candidate vaccine virus must be somewhat similar to current circulating strains, but not too similar (27). Quantifying the similarity of HA strains by correlating protein sequence with antigenicity can be achieved using antigenic cartography, which is a method for mapping the antigenic distance of related protein sequences in a two-dimensional space (10, 28). For rigorous quantification of antigenicity, we assert that the similarity of glycosylation state of the candidate virus HA to circulating strains must be included as well. However, a quantitative representation of glycosylation similarity between two strains is not a trivial task due to the heterogeneities of glycoforms and occupation state. Furthermore, because IAV utilizes host glycosylation machinery, the specific expression platform used to manufacture the vaccine greatly affects the resulting glycosylation of the vaccine virus.

Cao et al published a method for semi-quantification of viral glycoprotein glycosylation using differential glycosidase digestion and proteomics (29, 30). This method determines percent site occupancy and determines the degree to which a glycosite is occupied by high mannose or complex-type *N*-glycans. For comparison of glycosylation similarity, however, it is necessary to quantify each glycosite glycoform as rigorously as possible. Currently, there are no well-tested methods for making such rigorous glycosylation similarity comparisons among virus samples. The challenge is that the ability to both identify and quantify sample glycopeptide glycoforms is limited by mass spectrometer instrument speed.

Here, we present a method, using the Tanimoto similarity metric (31, 32), for the quantification of statistically rigorous similarity between two related virus strains that considers the presence and abundance of glycopeptide glycoforms. We describe the importance of liquid chromatography-mass spectrometry (LC-MS) data quality on the ability to make rigorous statistical similarity comparisons. Using the Tanimoto metric, we demonstrate using this approach to estimate similarity between two sources of IAV, both site-specifically and at the whole protein level.

## Experimental Procedures

### Generation of influenza viruses using reverse genetics

The full-length cDNA for HA or a mutated gene and neuraminidase genes of influenza A/Switzerland/9715293/2013 (H3N2) virus were amplified by using SuperScript One-Step RT-PCR (Invitrogen, Grand Island, NY), and the 6:2 recombinant viruses with six internal genes of influenza A/PR/8/1934(H1N1) virus were generated by using reverse genetics as described elsewhere (33). The mutated HA gene has three mutations Q132H, Y219S, and D225N, compared with the wild type HA gene, and we name the mutant as 5B8.

### Viral glycopeptide sample preparation

A/Switzerland/9715293/2013 (H3N2) virus and the mutant 5B8 virus were propagated in specific pathogen free embryonated chicken eggs at 37 °C for 72 hrs. A/Philippines/2/1982 (Phil82) samples, expressed in embryonated chicken eggs, were a generous gift from Dr. Kevan Hartshorn of Boston University School of Medicine. The viruses were purified using ultra-centrifuge as described elsewhere (34).

Viral samples were sonicated for 15-20 min in methanol to lyse virion membranes, and then evaporated to dryness in a centrifugal evaporator. Glycoprotein solutions were denatured and reduced with 100 mM ammonium bicarbonate (J.T. Baker), 2-2-2 trifluoroethanol (TFE) (Sigma, #T63002), and 200 mM dithiothreitol (DTT) (Sigma), incubating for 1 hr at 65 °C. Cysteine residues were alkylated by incubating with 200 mM iodoacetamide (Bio-Rad) for 1 hr at room temperature in the dark. After alkylation, excess iodoacetamide was quenched with additional DTT, again incubating for 1 hr at room temperature in the dark. The amount of TFE in each mixture was diluted to at most 5% with the addition of a 3:1 solution of water:100 mM ammonium bicarbonate. Sequencing-grade trypsin or chymotrypsin (Promega) was added at an enzyme:substrate ratio of 1:20, and the solutions were incubated overnight at 37 °C. After incubation, the enzymes were inactivated by heating the samples for 10 min at 95 °C. A small aliquot from each sample was reserved for de-glycosylation, and the remainders were enriched using iSPE-HILIC solid-phase extraction columns (HILICON), as follows. Samples were evaporated to dryness, and the HILIC columns were conditioned with water, then acetonitrile, and equilibrated with 80% acetonitrile/1% trifluoroacetic acid (TFA). Immediately prior to loading onto HILIC SPE columns, samples were reconstituted in 80% acetonitrile/1% TFA. Columns were washed in sample loading buffer, and then eluted with 1% acetonitrile/0.1% formic acid. Samples were evaporated to dryness, and then resuspended in 1% acetonitrile/0.1% formic acid for LC-MS injection.

The reserved aliquots were incubated with 500 units PNGase F (New England Biolabs) for every 20 μg of glycoprotein overnight at 37 °C. After incubation, de-glycosylated samples were cleaned using PepClean C18 spin columns, the eluate containing de-glycosylated peptides.

### Exoglycosidase modification of AGP and A/Philippines/2/1982 samples

Alpha-1-acid glycoprotein (AGP) purified from human serum (Sigma #G9885) and A/Philippines/2/1982 (Phil82) were digested in parallel with trypsin and chymotrypsin, as above. Before HILIC enrichment, the digested AGP glycopeptides were incubated with 200 units of α2-3,6,8 Neuraminidase (New England Biolabs # P0720S) for every 20 μg of glycoprotein overnight at 37 °C. This was to remove all terminal sialic acid residues from AGP glycans. The following day, the asialo AGP glycopeptides and digested Phil82 glycopeptides were divided into two equal aliquots. The first aliquot was HILIC-enriched, as above. The second aliquot was incubated overnight at 37 °C with β1-4 galactosidase (New England Biolabs #P0745) using the same reaction conditions as the neuraminidase above, and then HILIC-enriched. The aliquots with terminal galactose residues removed were denoted AGPgal and Phil82gal.

### LC-MS/MS acquisition

Liquid chromatography-tandem mass spectrometry (LC-MS/MS) data for de-glycosylated peptides were acquired in duplicate; those for HILIC-enriched glycopeptide samples were acquired with at least three replicate injections. All samples were analyzed by LC-MS/MS using a Q Exactive HF mass spectrometer (Thermo-Fisher) in positive mode, equipped with a nanoAcquity UPLC system, (nanoAcquity NPLC Symmetry C18 trap column and ACQUITY UPLC Peptide BEH C18 analytical column; Waters) and a Triversa Nanomate source (Advion, Ithaca, NY). Using mobile phase A (1% acetonitrile/0.1% formic acid) and mobile phase B (99% acetonitrile/0.1% formic acid), peptides were trapped for 4 min at 4 μL/min in A, and then separated using a flow rate of 0.5 μL/min with the following gradient: 0–3 min: 2–5% B, 3–93 min: 5–40% B, 93–98 min: 40% B, 98–100 min: 40–98% B, 100–105 min: 98% B, 105–106 min: 98–2% B, and 106–120 min: 2% B. Data for both de-glycosylated peptides and HILIC-enriched glycopeptides were acquired using data-dependent acquisition (DDA) methods. MS1 spectra were acquired at 60,000 resolution at *m/z* 400, scan range *m/z* 350–2000, 1 microscan per spectrum, AGC target of 1e6, maximum injection time 200 ms (100 ms for de-glycosylated peptides). MS2 spectra were acquired at 15,000 resolution at m/z 400, 2 microscans per spectrum, AGC target 3e6 (1e6 for de-glycosylated peptides), maximum injection time 200 ms, isolation window 2.0 m/z, isolation offset 0.4 *m/z*, exclusion of unassigned charge states and charge states 1, dynamic exclusion 10 s, and stepped NCE at 30, 40 (NCE 27 for de-glycosylated peptide). The 20 most abundant precursor ions per scan were fragmented. Profile data were recorded for both MS1 and MS2 scans.

## Data analysis

All raw files were first converted to mzML format (35) using MSConvert (36) with no additional filters.

Proteomics data from de-glycosylated samples were processed using the “Peaks PTM Search”function of the Peaks Studio 8.5 software (Bioinformatics Solutions, Waterloo, ON) (37). AGP was searched against the entire human Uniprot proteome (UP000005640 9606) (38). Databases for Phil82 and the SWZ13 variants consisted of HA and neuraminidase (NA) sequences from each specific strain and internal IAV protein sequences from A/Puerto Rico/8/1934 (PR8) appended to the host proteome, either *Gallus gallus* (UP000000539 9031) or *Canis lupus familiaris* (UP000002254 9615). Relevant FASTA files can be found in Figures S1 and S2. Technical replicates were searched together. Trypsin or chymotrypsin was specified as the proteolytic enzyme, with maximum of 3 missed cleavages and one non-specific cleavage were allowed. Cysteine carbamidomethylation and methionine oxidation were specified as fixed and variable modifications, respectively. The precursor ion (MS1) mass error tolerance was specified at 10 ppm and a fragment ion (MS/MS) error tolerance at 0.02 Da. Protein identifications required a minimum of two unique peptides. Default settings were used for setting the threshold score for accepting individual spectra. We used a target-decoy false discovery threshold of 1%. All peptides identified by Peaks Studio are listed in Supplemental File 1, including precursor charge, m/z, all modifications observed, peptide identification score, accession numbers, and protein groups of the proteins corresponding to the identified peptides. Protein abundances were calculated by aggregating the MS1 peak areas for all the peptides deconvoluted and identified for a specific protein; Supplemental File 2 shows the aggregated areas and the number of distinct peptides assigned for each protein. Peaks Studio search results were exported in mzIdentML (24) format. The mzIdentML files and all raw files have been deposited to the ProteomeXchange Consortium (http://proteomecentral.proteomexchange.org), via the PRIDE partner repository (36) with the dataset identifier PXDxxxxxx and DOI 10.6019/PXDxxxxxx.

LC/MS/MS raw files for all intact HILIC-enriched glycopeptides were processed using the GlycReSoft search engine, a complete open-source software package that assigns glycopeptides from tandem mass spectra (39). GlycReSoft requires the input of a peptide hypothesis, obtained in this work from the Peaks Studio proteomics search described above, and a glycan hypothesis. We constructed the glycan hypothesis combinatorially, following these glycan composition rules: #Hex 3-10, #HexNAc 2-9, #Fuc 0-5, #Neu5Ac 0-4, with #Fuc < #HexNAc and #HexNAc > #Neu5Ac + 1. The same glycan hypothesis was used for AGP and IAV samples, alike. GlycReSoft default values were used for modification localization thresholds. Glycopeptide abundances were quantified by aggregating the MS1 peak areas of the glycoforms deconvoluted and identified by GlycReSoft. All replicate runs for the glycopeptide assignments of a particular sample were combined into one table, displaying glycopeptide abundances for each replicate (Supplemental File 3). Tables and annotated spectra for all glycopeptides, including those with ambiguous localizations are provided in Supplemental Files 4-12.

### Bioinformatics similarity analysis

The glycopeptide identifications from the GlycReSoft searches included variations in peptide backbone resulting from missed cleavages, or variable oxidation of methionine or deamidation of asparagines. Before performing the bioinformatics analyses, the abundances of all peptide variants from a given sequon with the same glycan composition were summed together so that for each sequon, glycan compositions were not duplicated. In addition, glycoforms with only a single observation across replicates were discarded.

We used a modified Tanimoto coefficient, *T*, to calculate the similarity of glycosylation between two systems:

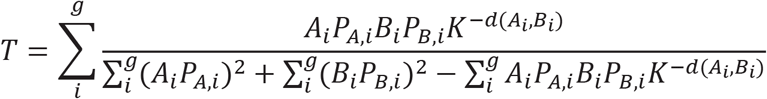

**A** and **B** are vectors assigned to the experimental groups being compared, containing the abundance values of each glycopeptide in the set. For each glycopeptide, the abundances are log scaled to give low weight to low abundance peaks, standardized to be in range [0,1], and then averaged across technical replicates, not including missing values. *P*_A_ and *P*_B_ are vectors containing the proportion of observed values to the total possible observations across technical replicates. *K* is a distance scaling term equal to [1+ *mean*(*P*_A_, *P*_B_)], and *d*(A,B) is the Manhattan distance between **A** and **B**. First, a “null” distribution composed of Tanimoto coefficients is calculated by comparing randomly selected combinations (with replacement) of replicates from **A** and **B** groups combined. The null distribution should be near 1 (very high similarity) because the groups are sampled equally. A “test” distribution is made by calculating the Tanimoto coefficients of randomly selected combinations of replicates from group **A** with those from group **B**. The observed Tanimoto similarity is the Tanimoto coefficient calculated from all replicates from **A** to all those of **B**. The degree of overlap of the null and test distributions reflects the confidence with which we can quantify the glycosylation similarity between the two groups.

### Experimental design and statistical rationale

For each enzyme-digested sample of AGP, AGPgal, Phil82, and Phil82gal, at least three technical replicates were acquired. Due to malfunction of the LC unit, one replicate for Phil82 chymo and mutant 5B8 IAV HA (chymo) were run on a different Aquity LC unit, but with the same mobile phases and method. As a result, we expected that there would be more differences in glycopeptide identifications and abundances for these two samples between replicates.

**Table 1.** Sample names and replicates.

| Sample type | Enzyme | # replicates |
| --- | --- | --- |
| WT SWZ13 HA<sup>a</sup> | Chymotrypsin | 3<sup>c</sup> |
|  | Trypsin | 4 |
| mutant 5B8HA<sup>b</sup> | Chymotrypsin | 3 |
|  | Trypsin | 3 |
| AGP | Chymotrypsin | 3 |
|  | Trypsin | 3 |
| AGPgal | Chymotrypsin | 3 |
|  | Trypsin | 3 |
| Phil82 | Chymotrypsin | 3<sup>c</sup> |
|  | Trypsin | 3 |
| Phil82gal | Chymotrypsin | 3 |
|  | Trypsin | 3 |

<sup>a</sup>Wild-type IAV strain A/Switzerland/9715293/2013, expressed in egg.
<sup>b</sup>Mutant IAV strain A/Switzerland/9715293/2013, expressed in egg.
<sup>c</sup>One replicate was run using a different LC unit, but with the same LC parameters.

AGP and Phil82 were used as control samples. All samples and technical replicates were acquired by LC-MS in randomized order.

## Results

### Tanimoto similarity coefficients

The Tanimoto similarity metric comparing AGP vs. AGPgal, Phil82 vs. Phil82gal, and mutant 5B8 vs. WT SWZ13 HA, were calculated for the proteins as a whole and site-specifically. Mutant 5B8 HA had three amino acid substitutions from the WT HA, Q132H, Y219S, and D225N. Similarity distributions are presented in Figure 2 and Table 2 for the whole protein comparisons, and in Figures 3, 4, and 5, and Table 3 for the site-specific comparisons.

**Table 2.** Tanimoto similarities for all whole-protein comparisons. Comparison groups are perfectly dissimilar when Tanimoto coefficient is 0 and perfectly similar when the coefficient is 1.

| Comparison | Enzyme | Tanimoto coefficient |
| --- | --- | --- |
| AGP vs. AGPgal | Chymotrypsin | < 0.0001 |
|  | Trypsin | < 0.01 |
| Phil82 vs. Phil82gal | Chymotrypsin | 0.65 |
|  | Trypsin | 0.62 |
| mutant 5B8 vs. WT<br>SWZ13 HA | Chymotrypsin | 0.79 |
|  | Trypsin | 0.77 |

**Table 3.**
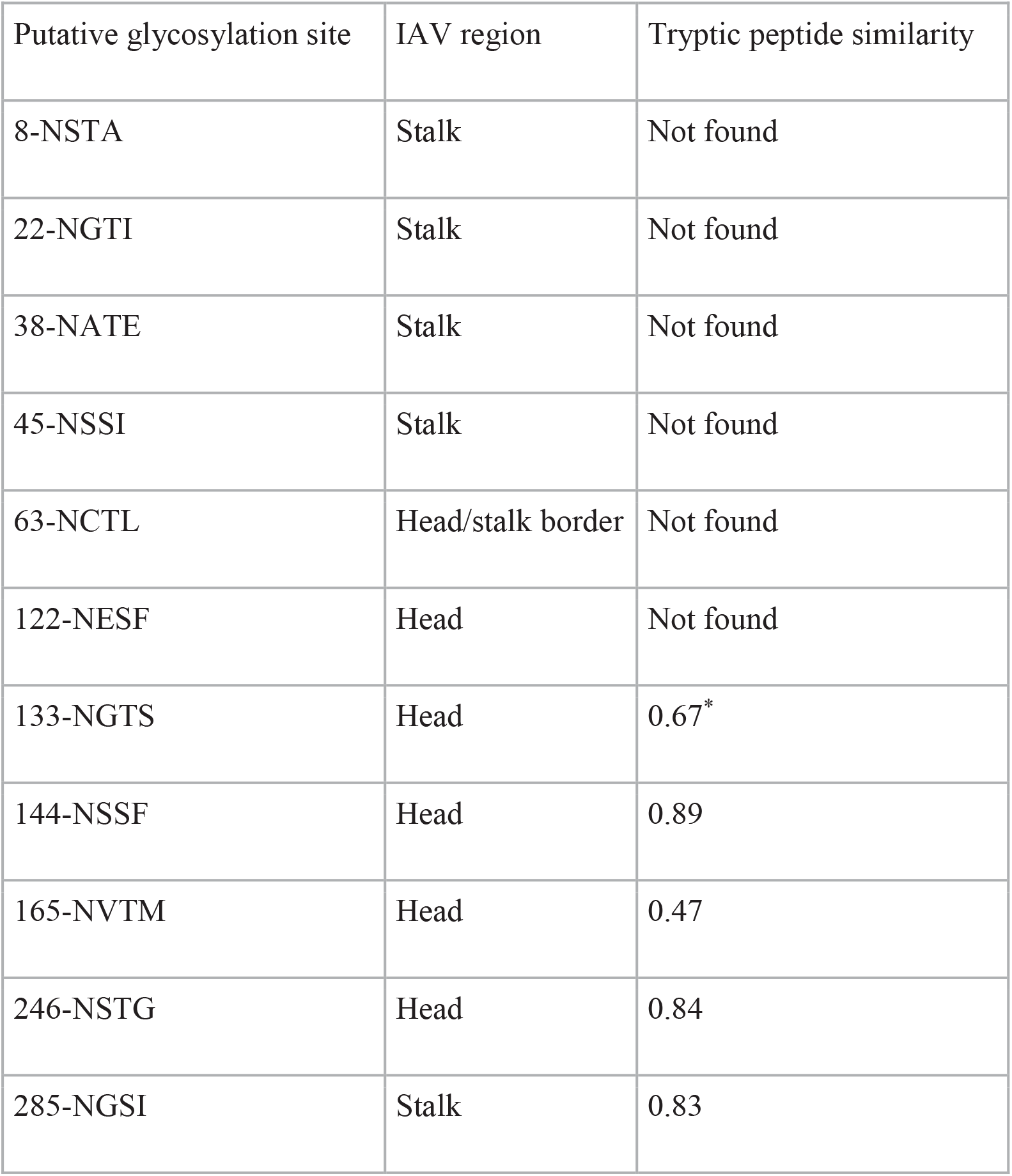
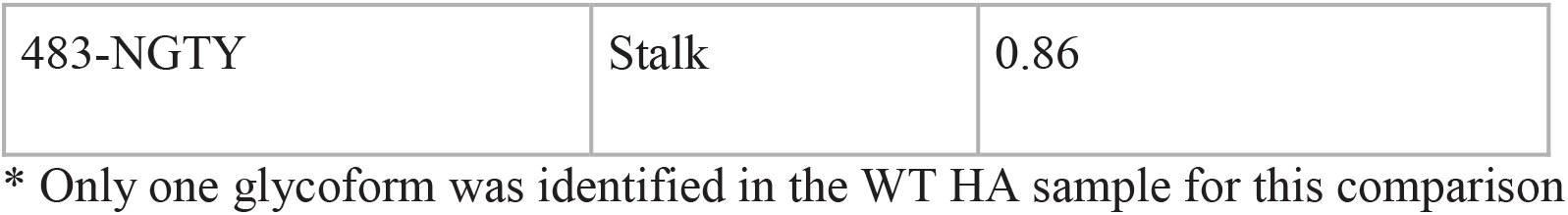
Site-specific similarity between mutant 5B8 and WT SWZ13 HA.

| Putative glycosylation site | IAV region | Tryptic peptide similarity |
| --- | --- | --- |
| 8-NSTA | Stalk | Not found |
| 22-NGTI | Stalk | Not found |
| 38-NATE | Stalk | Not found |
| 45-NSSI | Stalk | Not found |
| 63-NCTL | Head/stalk border | Not found |
| 122-NESF | Head | Not found |
| 133-NGTS | Head | 0.67* |
| 144-NSSF | Head | 0.89 |
| 165-NVTM | Head | 0.47 |
| 246-NSTG | Head | 0.84 |
| 285-NGSI | Stalk | 0.83 |
| 483-NGTY | Stalk | 0.86 |

\* Only one glycoform was identified in the WT HA sample for this comparison

### Tanimoto coefficient plots

We used the Tanimoto coefficient to estimate glycosylation similarity between two strains of HA, taking into account the abundances of the distribution of glycan compositions, including measurement variabilities. As shown in the example plot in Figure 1A, we represented the data visually by plotting null and test distributions of glycans on a similarity scale. The x-axis shows the Tanimoto similarity. The y-axis is the density of the distributions (analogous to frequency in a histogram plot).

**Figure 1.**
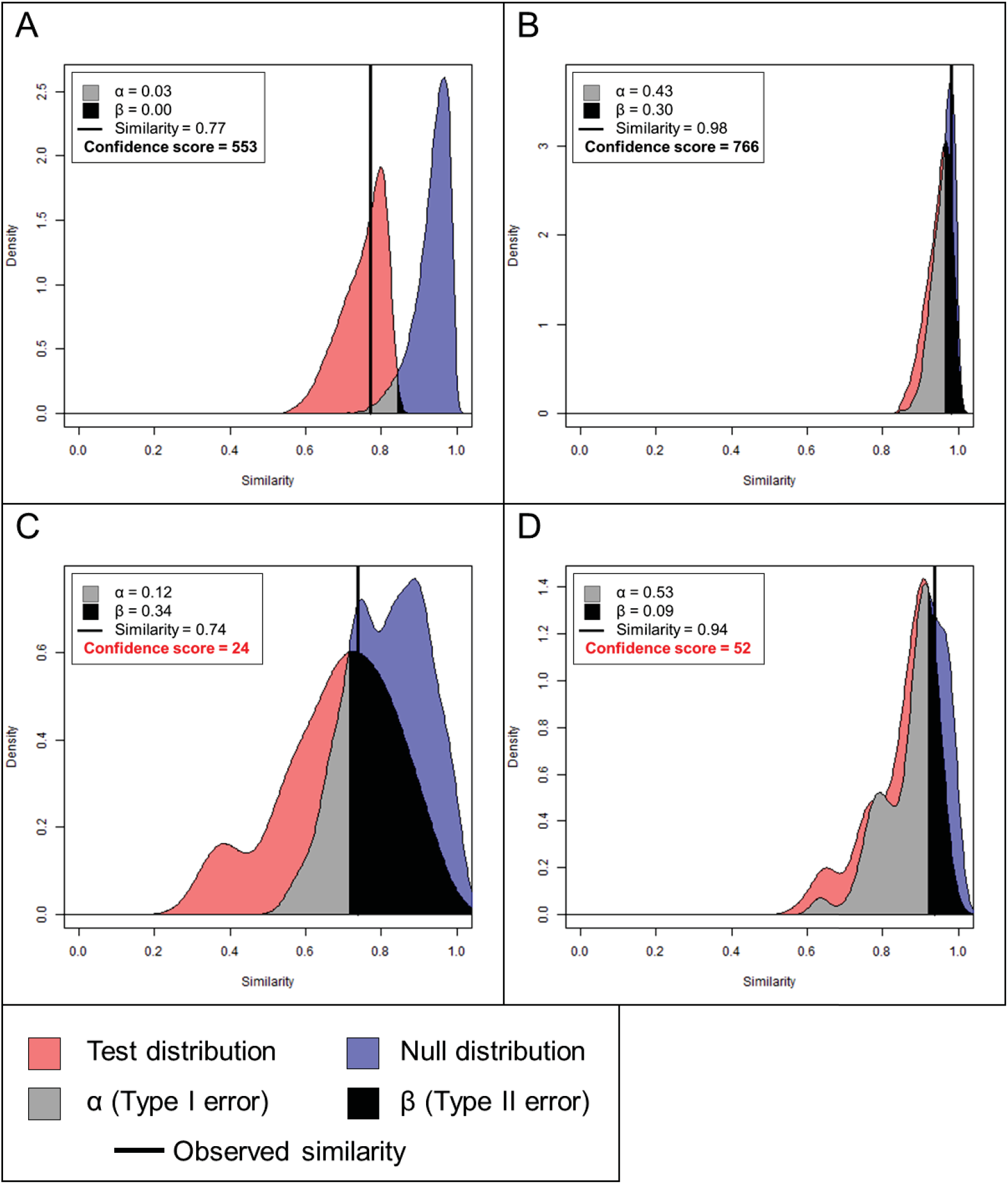
Example Tanimoto plots. Tanimoto similarity is represented on the x-axis, with perfectly dissimilar at 0 and perfectly similar at 1. The observed similarity is represented by the vertical black line. The α and β values are the Type I and II errors, respectively. A confidence score of ≥77 indicates good confidence. (A) Ideal plot with narrow null and test distributions and good resolution between the two. (B) Poorly resolved null and test distributions, but the distributions are narrow and the confidence score is high, indicating that the two groups are highly similar. (C) Poorly resolved null and test distributions, but distributions are wide, indicating poor data quality, and the confidence score is < 77. (D) Inconsistencies among sample replicates results in multimodal distributions.

The null distribution (blue area in Figure 1A) was made by calculating the Tanimoto coefficient between randomly selected combinations (with replacement) of replicates from sample 1 and sample 2 combined. Because these combinations of replicates were randomly selected from the same combined dataset, the distribution of Tanimoto coefficients should be close to 1, or perfectly similar. The test distribution (red area), was composed of Tanimoto coefficients comparing randomly selected combinations (with replacement) of replicates from sample 1 to those of sample 2. The observed Tanimoto similarity (the thick vertical line) is the Tanimoto coefficient calculated from all the replicates of sample 1 to all those of sample 2.

The null and test distributions and their overlap are analogous to null and alternative hypotheses and the test statistic in statistical inference. That is, if the null and test distributions do not overlap significantly, we then reject the null hypothesis and conclude that the two distributions are significantly dissimilar. Else, we fail to reject the null hypothesis because we do not have statistically significant evidence to conclude that the distributions are dissimilar.

The area of overlap to the left of the point of intersection between the two distributions (grey area) is analogous to the Type I Error (α), or the false positive rate, and the area to the right (black area), the Type II Error (β), or false negative rate (equivalently, 1 – power). A significance level of α = 0.05 (i.e. the grey area is 5% of the joint area of both the null and test distributions) means that for 5% of Tanimoto coefficients in the joint test and null distributions, we do not have statistically significant evidence to determine whether they are false positives or not, and therefore, we conservatively conclude they are false positive. A power of 80% (i.e. β = 0.2) means that the black area is equal to 20% of the joint area of the null and test distributions. A black overlap area of greater than 20% indicates that our data are not sufficiently powered to make a confident conclusion about the similarity between samples.

Figure 1A is an example of an ideal plot with narrow null and test distributions and good resolution between the two. The overlap areas were α = 0.03 and β = 0.00, indicating acceptable levels of Type I and II errors, respectively. Figure 1B-D are examples of poorly resolved distributions. The distributions in Figure 1B overlapped substantially (α = 0.43 and β = 0.30; however, each individual distribution is narrow, indicating that these samples are actually highly similar. Figure 1C had very wide distributions as a result of large variances within both distributions. This may be an indication of poor data quality, and without further experimental validation, we cannot conclude that these samples are dissimilar. Figure 1D is an example of samples that have one or more replicates that were very distinct from the others, resulting in clustering of some comparisons within the distribution that shows up in the plot as multimodal distributions, an indication that there may have been problems with some sample replicates.

As a data quality control measure, we can overlay internal distributions, composed of Tanimoto coefficients from randomly selected combinations of replicates from a single sample, over the null and test distributions. The plots in Figure S3 are the internal quality plots corresponding to the four plots in Figure 1. For both samples 1 and 2 in Figure S3A, the internal distributions were tall and narrow, an indication of good data quality. We can measure the tallness and narrowness of the internal distribution by calculating the ratio of height to variance. In Figure S3A, the distribution for sample A had a height of 4.37 and a variance of 0.00054. Therefore, the distribution ratio for sample 1 was 8135; that for sample 2 was 570. For our purposes, an acceptable internal distribution ratio is ≥ 100, that is, the distribution should be at least 100 times as tall as it is wide to be considered good quality. All internal distribution ratio information can be found in Table S1.

Our confidence of the observed similarity depends on the α and β overlap areas in addition to the internal distribution ratios. For our data, we combined these pieces of information to obtain one confidence score, defined as *e*^−(α+β)^ multiplied by the lower of the two internal distribution ratios. The minimum acceptable confidence would be when α + β = 0.25, a minimum internal ratio of 100, or equivalently, *e*^−0.25^ × 100 = 77. The example plots in Figure 1A and 1B both had scores greater than 77. Those in Figure 1C and 1D were both less than 77, and therefore, we do not have enough evidence to make a conclusion about their similarities.

### Similarity of exoglycosidase-generated pairs

To evaluate the performance of the Tanimoto coefficient, we used well-characterized standard glycoproteins including AGP. The A1AG1 isoform of AGP has five *N*-glycosylation sites, each containing a distribution of complex glycans (40). If we modify those glycans by removing all terminal sialic acid and galactose residues, then the glycosylation of the modified protein, which we denote as AGPgal, is expected to have zero similarity to the unmodified protein. This turned out to be the case (Figure 2A and 2B). Because AGP has only complex glycans, nearly all were modified by the combination of sialidase and galactosidase. The Tanimoto distributions for both tryptic and chymotryptic digestions for the A1AG1 protein as a whole (Figure 2A and 2B) showed a similarity of < 0.01 and < 0.0001 with confidence scores of 466,987 and 6146, respectively. Site-specific distributions for chymotryptic and tryptic digestions are shown in Figure 3 and S6, respectively. Bar plots showing the standardized mean abundance for glycoforms at each sequon are also shown in these figures. Site 56-NKSV has low coverage in a tryptic digestion compared to the chymotryptic digestion. We identified only three tryptic 56-NKSV glycoforms from AGP and AGPgal, with the result that the null distribution for this comparison was very broad, as too were the internal distributions from the internal quality plots (Figure S6), a confirmation that the data quality was poor for this site. The confidence score for tryptic 56-NKSV was 6.

**Figure 2.**
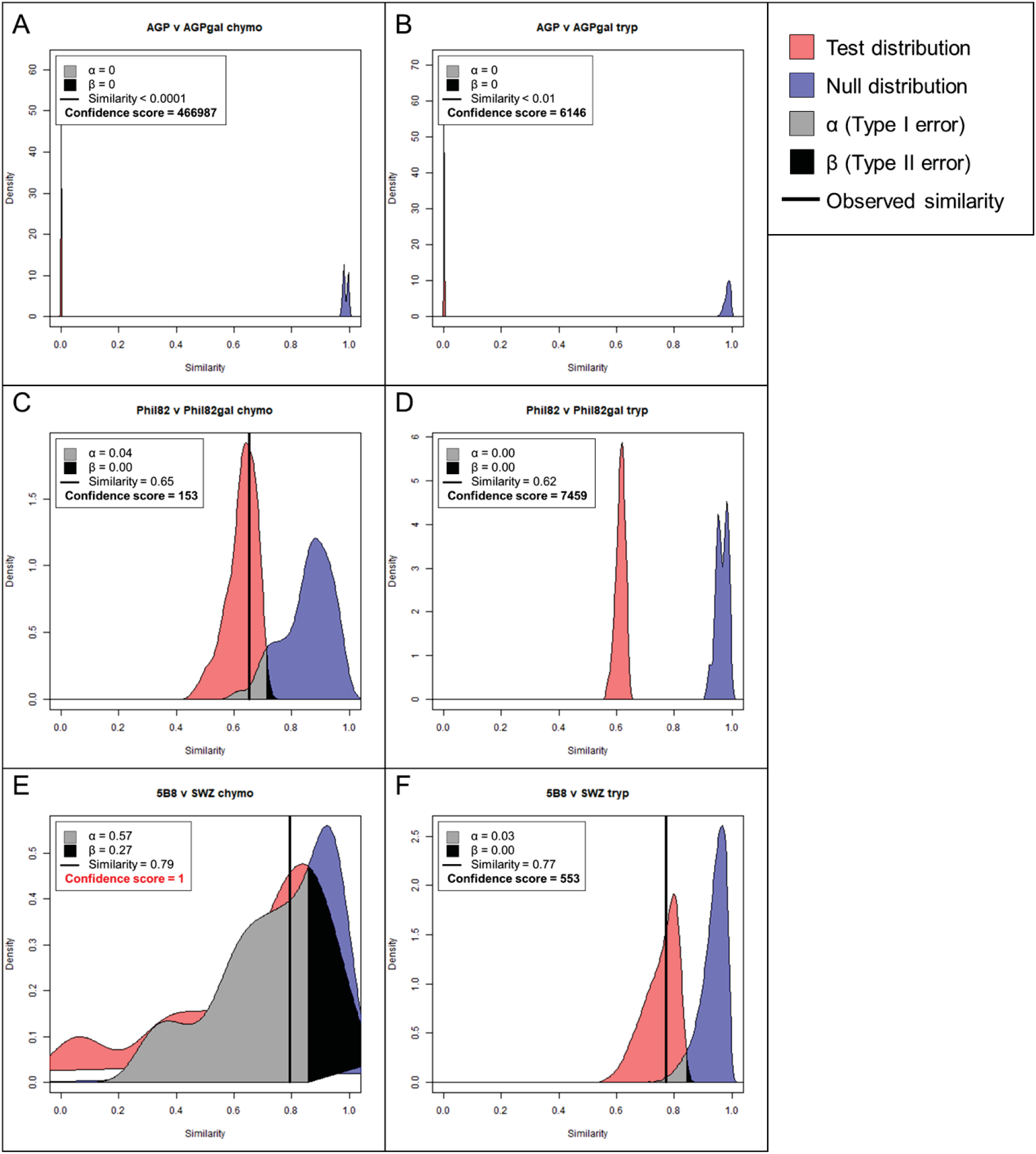
Tanimoto similarity at the whole protein level between (A) chymotryptic AGP and AGPgal, (B) tryptic AGP vs. AGPgal, (C) chymotryptic Phil82 HA vs. Phil82gal HA, (D) tryptic Phil82 HA vs. Phil82gal HA, (E) chymotryptic mutant 5B8 HA vs. WT SWZ13 HA, and (F) tryptic mutant 5B8 vs. WT SWZ13 HA. The observed similarity is represented by the vertical black line, α and β are the Type I and II errors, respectively. A confidence score of ≥77 indicates good confidence. Tanimoto similarity is represented on the x-axis, with perfectly dissimilar at 0 and perfectly similar at 1.

**Figure 3.**
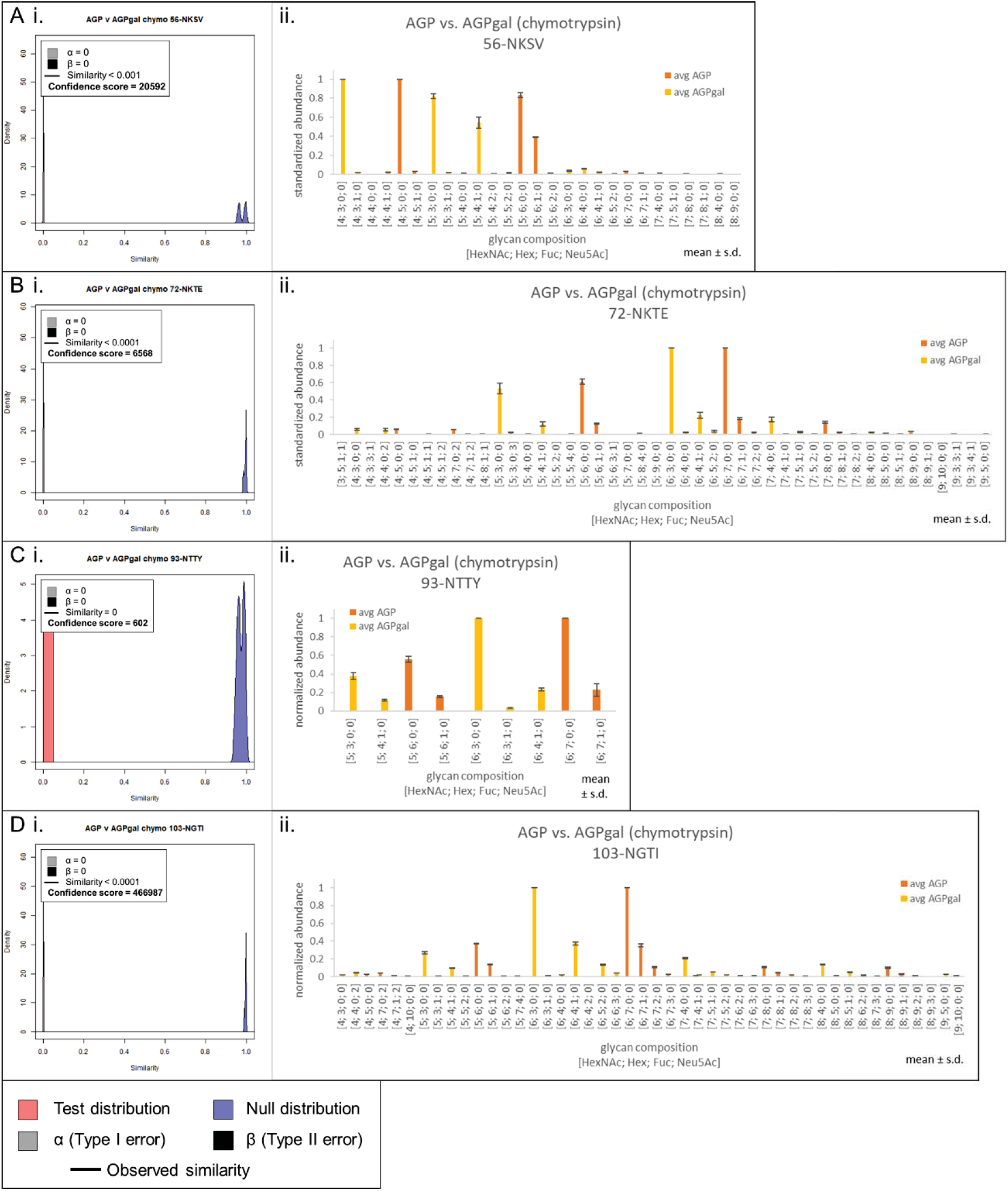
Site-specific comparisons for chymotryptic glycopeptides of AGP and AGPgal. Site 33-NATL was not found for chymotrypsin. (A) Tanimoto distribution plot (i) and standardized abundance bar plots (ii) for AGP vs. AGPgal at site 56-NKSV, (B) site 72-NKTE, (C) site 93-NTTY, and (D) site 103-NGTI. In the Tanimoto distribution plots, the observed similarity is represented by the vertical black line, α and β are the Type I and II errors, respectively. A confidence score of ≥77 indicates good confidence. Tanimoto similarity is represented on the x-axis, with perfectly dissimilar at 0 and perfectly similar at 1. In the bar plots, the glycopeptide abundances were standardized to be in range [0,1], and averaged across technical replicates, not including missing values. Error bars show ± standard deviation.

A/Philippines/2/1982 (Phil82) is a more complex and biologically relevant example for which we demonstrate the robustness of the Tanimoto metric. IAVglycans are largely asialo because the neuraminidase (NA) envelope protein cleaves sialic acid residues so that newly formed virions can bud off from the host cell’s plasma membrane. Again, we modified the Phil82 HA glycans with β-galactosidase to remove any terminal galactose residues; we denote this sample as Phil82gal. We know from our previous work on site-specific characterization of Phil82 (24) that some Phil82 glycosites have primarily high-mannose *N*-glycans. These high-mannose glycans would not be cleaved, and therefore, the sites with high-mannose glycans would retain some degree of similarity before and after β-galactosidase treatment, depending on the abundance of the high-mannose glycans. By contrast, Phil82 glycosites occupied by complex *N*-glycans that have terminal galactose residues would be cleaved and show low similarity after cleavage by β-galactosidase. The Tanimoto plot comparing trypsin-digested Phil82 and Phil82gal data sets for the HA protein as a whole showed a moderately high similarity of 0.62, with confidence score 7459 (Figure 2D). However, the chymotryptic comparison showed a wide null distribution, which suggests data quality issues. In this particular case, due to poor LC performance, we were not able to acquire three replicates for chymotryptic Phil82gal using the same LC unit. Because our Tanimoto algorithm required a minimum of three replicates, the third technical replicate was acquired using a different LC unit. Although the confidence score was still above 100 (Figure 2C, we opted to discard the entire Phil82 vs. Phil82gal chymotryptic dataset because of the known inconsistency of LC in the data collection. Consequently, we performed all subsequent analyses using only the tryptic dataset. Total ion chromatograms for this and all other comparisons were made using Thermo Xcalibur Qual Browse 2.2 and are shown in Figures S8, S9, and S10.

Similarities for the six glycosylation sites that were observed in the tryptic Phil82 vs. Phil82gal comparison and their confidence scores can be found in Figure 4, along with bar plots comparing the standardized mean abundance for all identified glycoforms. These comparisons reflected the proportion of high-mannose *N*-glycans that occupy these sites. Sites 144-NNSF and 483-NGTY had high proportions of complex *N*-glycans in Phil82, and the similarities to their corresponding sites in Phil82gal were < 0.01 and 0.03, respectively. Sites 165-NVTM and 285-NGSI had high proportions of high-mannose *N*-glycans, and consequently, the similarities were high for these two sites, at 0.98 and 0.76, respectively. Site 38-NATE, having a mixture of both high-mannose and complex *N*-glycans, had a similarity in the middle at 0.69. Site 246-NSTG had high-mannose glycans and an observed similarity of 0.70, but poor data quality resulted in very broad distributions. As shown in Figure S11, the internal quality plot for Phil82 at the same site could not be made because there was only one glycoform identified in the Phil82 sample for this site, and as a result, the confidence score for this comparison was 0. Additional experimental validation is needed to make a confident assessment of the similarity at this site.

**Figure 4.**
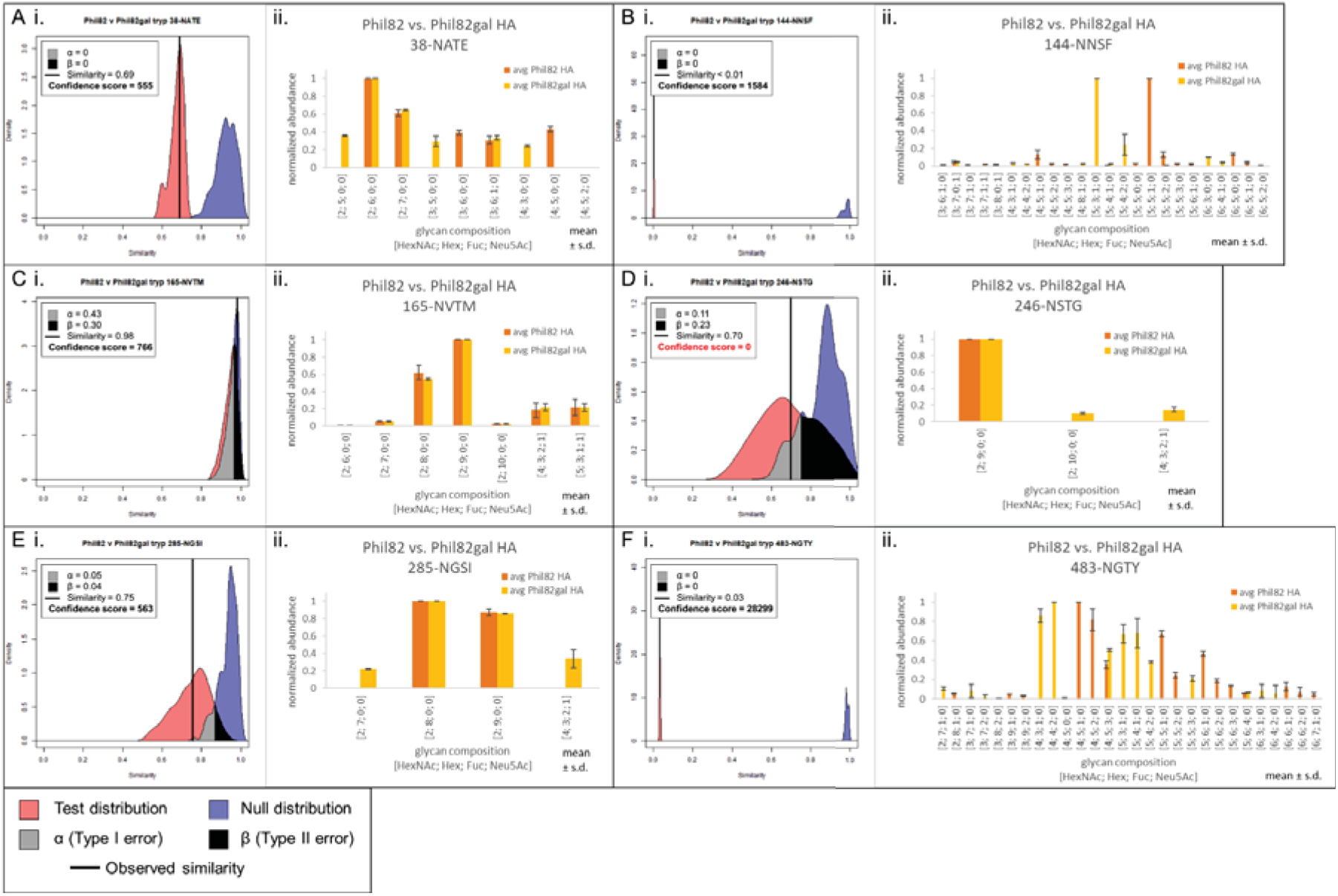
Site-specific comparisons for tryptic glycopeptides of Phil82 HA and Phil82gal HA. (A) Tanimoto distribution plot (i) and standardized abundance bar plots (ii) for site 38-NATE, (B) site 144-NNSF, (C) site 165-NVTM, (D) site 246-NSTG, (E) site 285-NGSI, and (F) site 483-NGTY. In the Tanimoto distribution plots, the observed similarity is represented by the vertical black line, α and β are the Type I and II errors, respectively. A confidence score of ≥77 indicates good confidence. Tanimoto similarity is represented on the x-axis, with perfectly dissimilar at 0 and perfectly similar at 1. In the bar plots, the glycopeptide abundances were standardized to be in range [0,1], and averaged across technical replicates, not including missing values. Error bars show ± standard deviation.

### Similarity between IAV pairs

A/Switzerland/9715293/2013 (SWZ13) is a highly-glycosylated H3N2 strain used in the 2015-2016 influenza vaccine (41, 42). It has 12 putative *N*-glycosylation sites. We examined the WT SWZ13 and a mutant (denoted 5B8), both expressed in embryonated chicken eggs, for this paper. Mutant 5B8 HA has three amino acid substitutions from the WT, Q132H, Y219S, and D225N. None of these substitutions adds or deletes a glycosylation site. In order to address the question of the extent to which these changes affect the conformation of the HA trimer in changing the accessibility of glycan-processing enzymes, we compared the Tanimoto similarity plots for trypsin and chymotrypsin digested virus, respectively. However, like the chymotryptic Phil82 data discussed above, one replicate from the chymotryptic WT SWZ13 HA set was run on a different LC unit, causing data quality issues. The null and test distributions for this comparison overlapped almost completely and were multimodal, indicating clustering of replicates (Figures 2C and S3C). Therefore, only the tryptic comparisons will be further discussed. Taking together all observed glycosites for the trypsin-digested mutant 5B8 vs. WT HA comparison, the Tanimoto similarity between the two was relatively high with a value of 0.77 (Figure 2F.) The confidence score was 553, with well-resolved null and test distributions as well as sharp internal distributions, demonstrating that even changes that do not change any of the glycosylation sites can still result in a measurably distinct population of glycans and glycan abundances throughout the protein.

We examined which particular sites contributed most to the overall similarity between mutant 5B8 and WT SWZ13 HA glycoprotein. Among the individual glycosites and the corresponding internal quality plots (Figures 5 and S12), 133-NGTS had a confidence score of 0 because there was only one glycoform identified for WT HA and the internal quality plot could not be made for this site sample. For all other sites, internal quality plots and confidence scores indicated sufficient data quality, and the null and test distributions were well resolved, with the exception of 165-NVTM. The low confidence score for 165-NVTM was due to the broad internal distribution for mutant 5B8 HA (ratio = 48); however, the overlap between the test and null distributions was 0. For this site, therefore, we conclude that although the data quality can be better, the near-perfect resolution between the null and test distributions means that we can be confident in this observed similarity. Four sites had relatively high similarity (144-NSSF, 246-NSTG, 285-NGSI and 483-NGTY), while 165-NVTM had relatively low similarity (Table 3).

**Figure 5.**
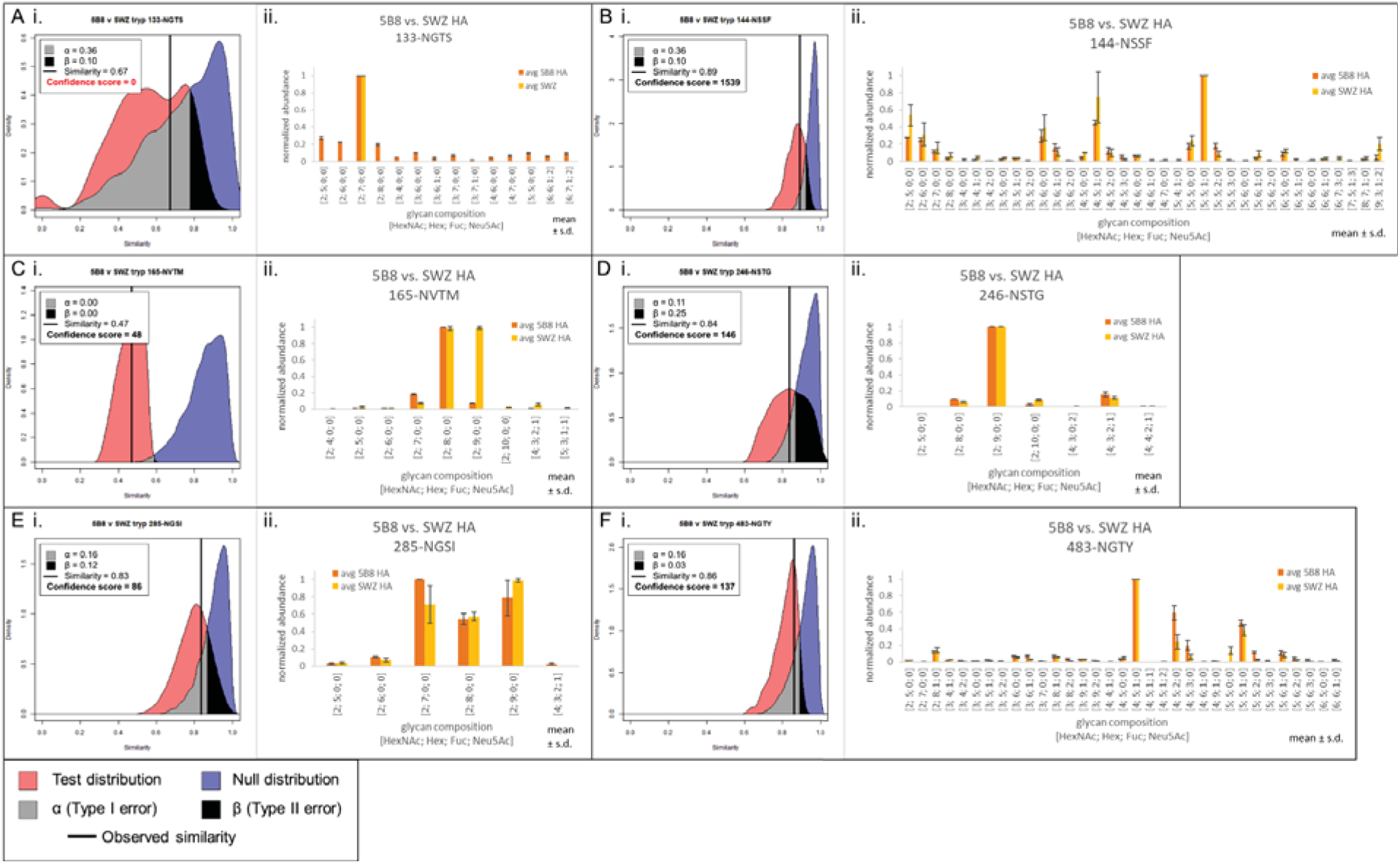
Site-specific comparisons for tryptic glycopeptides of mutant 5B8 HA and WT SWZ13 HA. (A) Tanimoto distribution plot (i) and standardized abundance bar plots (ii) for site 133-NGTS, (B) site 144-NSSF, (C) site 165-NVTM, (D) site 246-NSTG, (E) site 285-NGSI, and (F) site 483-NGTY. In the Tanimoto distribution plots, the observed similarity is represented by the vertical black line, α and β are the Type I and II errors, respectively. A confidence score of ≥77 indicates good confidence. Tanimoto similarity is represented on the x-axis, with perfectly dissimilar at 0 and perfectly similar at 1. In the bar plots, the glycopeptide abundances were standardized to be in range [0,1], and averaged across technical replicates, not including missing values. Error bars show ± standard deviation.

These data confirmed our expectation that the glycosylation site on the stalk region of HA (285-NGSI and 483-NGTY) exhibit a higher degree of glycosylation similarity and those on the globular head domain of HA (144-NSSF, 165-NVTM, and 246-NSTG) to exhibit variable similarity because the head receives more immune pressure. Antibodies specific to the antigenic regions of the head domain drive rapid genetic mutation, amino acid substitution, accumulation of new sequons in those regions, while the stalk remains relatively stable (43). We measured a similarity of 0.83 and 0.86 at 285-NGSI and 483-NGTY, respectively, both in the stalk region, which confirmed our expectations. However, the three sites in the head domain present a mixed range of similarities: 144-NSSF and 246-NSTG have high similarity (0.89 and 0.84, respectively), while 165-NVTM has a similarity of 0.47. Receptor binding sites are also located on the head region, and these sites are conserved across all strains of H3N2 IAV to preserve viral recognition of host receptors. These three head domain glycosylation sites are all located within or near to the receptor binding site and known antigenic sites (43), which means that there is competing pressure at these sites to retain the ability to recognize host cell receptors while promoting variation of its amino acid sequence to evade host cell immunity. Our data suggest that these competing pressures are reflected in the degree of similarity of glycosylation, in addition to whether the amino acid sequence changes or not. We observed that 144-NSSF and 246-NSTG had high glycosylation similarity, which suggests that these sites may be contributing to the architectural stability of the receptor-binding site, while 165-NVTM, which showed low glycosylation similarity between our two strains, may be exposed to antibody pressure. 165-NVTM has also been shown to bind to surfactant protein D, a collectin molecule that is part of the innate immune system (24, 44). Exposure to innate immune pressure may also be driving glycosylation variation at this site. As shown in the barplot for this site in Figure 5Cii, the main difference that contributes most to the low similarity was in the abundances of the Man8GlcNAc2 and Man9GlcNAc2 glycoforms. The mutant 5B8 HA has lower abundance of Man9GlcNAc2 compared to WT, and higher of Man8GlcNAc2, suggesting that the 5B8 mutations changed the conformation of the HA trimer enough to increase the accessibility of the glycan processing enzyme ER class I α-mannosidase (ERManI) (24, 45).

Although we cannot comment further on the chymotryptic results due to inconsistency between replicates, as discussed above, it is worth noting that the chymotrypsin digestion resulted in identification of glycosylation sites 8-NSTA and 38-NATE in addition to the six that were observed with trypsin. In future experiments, it may be worth considering that certain glycosylation sites are only accessible by chymotrypsin digestion

#### Improving acquisition methods

We explored various mass spectrometry methods tailored for glycoproteomics to achieve the highest reproducibility of data to inform most confidence in similarity calculations. Higher energy collisional dissociation (HCD) suffices for assignment of singly-glycosylated peptides (39). TopN data-dependent acquisition (DDA) is a common strategy for analysis of peptides. While this strategy is good for discovery of glycopeptide glycoforms, the stochastic nature of precursor selection by intensity results in replicates with a high number of missing values, limiting our ability to calculate similarity with high confidence. Furthermore, because glycopeptides are low in abundance compared to the unglycosylated peptides in the mixture, glycopeptide precursors often go unselected, further contributing to the missing value problem. Low reproducibility of DDA, therefore, necessitates exploring other acquisition options. We attempted parallel reaction monitoring (PRM) to maximize technical reproducibility, albeit at the expense of limited coverage of glycopeptide precursor ions in a given time window. PRM precursor ion selection is strictly deterministic. Precursors selected for fragmentation are programmed by the user prior to acquisition and are the same for every replicate of the sample, unlike DDA whose precursors are selected stochastically by intensity. The quantification for PRM is done at the MS2 level, using highly informative peptide-Y fragment ion series, imbuing higher confidence to the assignments and to the quantification. However, the capability to discover previously unidentified glycoforms is lost, and there is a limit to how many PRM targets can be selected due to the duty cycle of the mass spectrometer. As a result, fewer glycopeptides were subjected to tandem MS using PRM than DDA, resulting in unsatisfactory ability to compare complete glycosite glycoform distributions among biological samples, which is why we relied on aggregating deconvoluted MS1 precursor ions to determine glycopeptide abundances for this paper.

## Conclusions

Because the composition of the distribution of glycans that occupy a glycosite cannot be predicted from genomic data, quantification of glycopeptides from MS2 fragment ions is not straightforward and requires specialized bioinformatics tools. Quantitative proteomics methods including Skyline (31) allow a range of fixed chemical modifications to proteins, including phosphorylation, acetylation, methylation and acylation, among others(47–51). The landscape for quantitative glycoproteomics software, which must be able to handle a distribution of glycoforms at each glycosite, is far more limited.

We show that rigorous comparison of glycosite similarity requires careful consideration of glycoproteome complexity. The use of a measured glycome is considered preferable as a means of minimizing the number of glycan compositions used to calculate glycopeptide null distributions relative to use of combinatorial lists or entire glycan databases. Data confidence would be highest if it were possible to quantify all glycopeptide compositions using tandem MS. However, both DDA and PRM lack the speed necessary to quantify all such glycopeptide compositions for moderately complex glycoproteins such as HA, meaning that it is necessary to use MS1 peak abundances for which tandem MS confirmation of identity is not available. As the fraction of glycopeptides identified from MS1 peaks lacking supporting MS2 identifications increases, the overall confidence in the similarity decreases. We are therefore interested in the development of data-independent acquisition (DIA) methods for glycopeptides, which would in theory maximize reproducibility while allowing for the discovery of unknown glycopeptides, and providing more accurate quantification. In principle, the use of DIA for moderately complex glycoproteins such as HA would provide an advantage for similarity comparisons only if there were sufficient ability to select among the several related glycopeptides that elute in a narrow chromatographic retention time window. This is a topic for future work.

## Abbreviations

IAV: influenza A virus
HA: hemagglutinin
NA: neuraminidase
AGP: alpha-1-acid glycoprotein
AGPgal: asialo AGP with terminal galactose residues removed
Phil82: A/Philippines/2/1982
Phil82gal: Phil82 with terminal galactose residues removed
SWZ13: A/Switzerland/9715293/2013
5B8: SWZ13 with amino acid substitutions Q132H, Y219S, and D225N in HA

## Acknowledgements

This work was supported by NIH grants U01CA221234 and R01GM133963. LZ and XFW were supported by NIH grant 1R01AI116744.

## Data availability

The mass spectrometry data have been deposited to the ProteomXchange Consortium (http://proteomecentral.proteomexchange.org) via the PRIDE partner repository with the dataset identified PXDxxxxxx and DOI 10.6019/PXDxxxxxx.

